# Engineered membranes for residual cell trapping on microfluidic blood plasma separation systems. A comparison between porous and nanofibrous membranes

**DOI:** 10.1101/2020.05.21.107813

**Authors:** Francesco Lopresti, Ieva Keraite, Alfredo E. Ongaro, Nicola M. Howarth, Vincenzo La Carrubba, Maïwenn Kersaudy-Kerhoas

## Abstract

Blood-based clinical diagnostics require challenging limit-of-detection for low abundance, circulating molecules in plasma. Micro-scale blood plasma separation (BPS) has achieved remarkable results in terms of plasma yield or purity, but rarely achieving both at the same time. Here, we proposed the first use of electrospun polylactic-acid (PLA) membranes as filters to remove residual cell population from continuous hydrodynamic-BPS devices. The membranes hydrophilicity was improved by adopting a wet chemistry approach via surface aminolysis as demonstrated through Fourier Transform Infrared Spectroscopy and Water Contact Angle analysis. The usability of PLA-membranes was assessed through degradation measurements at extreme pH values. Plasma purity and hemolysis were evaluated on plasma samples with residual red blood cell content (1, 3, 5% hematocrit) corresponding to output from existing hydrodynamic BPS systems. Commercially available membranes for BPS were used as benchmark. Results highlighted that the electrospun membranes are suitable for downstream residual cell removal from blood, permitting the collection of up to 2 mL of pure and low-hemolyzed plasma. Fluorometric DNA quantification revealed that electrospun membranes did not significantly affect the concentration of circulating DNA. PLA-based electrospun membranes can be combined with hydrodynamic BPS in order to achieve high volume plasma separation at over 99% plasma purity.

## 1 Introduction

Blood plasma separation is usually required for several blood-based clinical diagnostics [1]. In this context, plasma purity superior to 99% is a crucial requirement since presence of blood cells and hemoglobin, may interfere with immunoassays and other assays [1–3]. On the other hand, cell lysis, such as hemolysis, but also leukolysis, should be avoided as DNA or RNA released from lysed cells may interfere with circulating cell-free DNA and RNA levels, resulting in erroneous readings, particularly in the case of liquid biopsy assays. In recent years, there has been a growing interest in micro-scale blood plasma separation (BPS) both as a replacement for traditional centrifugation and as preliminary modules to complete Lab-On-Chip devices. BPS studies have demonstrated remarkable results in terms of yield or purity, but have rarely achieved both at the same time [2].

Micro-scale BPS strategies adopted until now depend on several factors such as the volume of plasma required for subsequent clinical tests as well as the number and type of tests to be performed on the same sample. For instance, blood gas, electrolytes and metabolites (BGEM) tests can be carried out on plasma from a drop of blood, typically in the order of 10-200 µL. In this volume range, paper-based technologies or filtration are the most common approaches [4–7].

If larger blood volumes are needed (milliliter range), such in the case of amplification of rare circulating DNA from plasma [8], hydrodynamic blood plasma separation (HBPS) is preferred because it avoids the saturation usually quickly reached by membrane-based separation. These approaches permit achieving plasma purity close to 100%, although plasma yields are usually low (1-15%) and often the plasma purity decreases upon increasing plasma yields [9–13].

The combination of hydrodynamic and membrane separation in a BPS system is proposed to achieve high yield and high purity simultaneously, when processing large volumes of blood, by allowing large amount of plasma to be removed with suboptimal purity, followed by removal of residual RBCs by filtration without loss of plasma volume.

Membrane filtration at microscale is often used as a sieving process to retain large molecules or particulate matter [14–16]

However, in order to accomplish high performance separation, membranes should meet challenging requirements such as low pressure drop and high plasma purity. The development of engineered membranes could allow the control of chemical, physical and mechanical properties of biopolymeric highly porous structure susceptible to meet these requirements. Among the several approaches developed for fabricating biopolymeric porous structures, thermally or diffusion induced phase separation (TIPS or DIPS) as well as electrospinning are gaining more attention [17,18].

TIPS and DIPS techniques are based on thermodynamic de-mixing of a homogeneous polymer-solvent solution into a polymer-rich and a polymer-lean phase. De-mixing can be achieved by a change in temperature (TIPS) or by changing the exposure of the solution to another immiscible solvent (DIPS).[17,18] Both these approaches lead to the production of well interconnected porous structures exhibiting tunable pore size.

On the other hand, electrospinning produces ultrafine fibers ranging from micro-to nano-meter in diameters from polymeric solutions or melts [19]. Electrospun (ES) membranes show high porosity and, tunable mechanical properties as well as high surface to volume ratio due to their small fiber dimension and they were successfully integrated in microfluidic chips for cell culture or for immunoassay optimization [20–23].

One of the most common biopolymers used for these applications is polylactic acid (PLA). Due to its commercial availability at an affordable cost, PLA has been extensively studied and used in many biomedical purposes including wound healing [24,25] and tissue engineering scaffolds [26,27]. Recently, we have shown that PLA can be used as a novel substrates for microfluidic devices [28]. During the last decades, researchers have contributed to widening the application range of PLA through chemical and physical alterations to the bulk and/or surface by introducing hydrophilic and biocompatible moieties [29,30]. Several modifications strategies such as plasma [31] or chemical [29] treatments have been employed to enable specific surface properties on PLA. For instance, surface modification such as the aminolysis technique has been widely used for the addition of amine groups in order to covalently bind bioactive molecule on PLA [32].

In an effort to develop a rapid and effective residual cell trapping, the efficacy of PLA-based membranes prepared by TIPS, DIPS, and electrospinning for blood plasma separation was assessed. To the best of our knowledge, it is the first time that PLA-based porous membranes have been tested as a microfiltration unit (MFU) for microfluidic blood plasma separation purposes. Initially, a comparative study on PLA membranes degradation at extreme pH values was carried out in order to ascertain their usability in microfluidic protocols requiring the use of non-neutral buffers or after acid or basic treatments. Secondly, plasma purity, free hemoglobin concentration and pressure drop across the DIPS, TIPS and ES membranes have been evaluated employing plasma-diluted blood corresponding to output-plasma tested from already existing HBPS systems [33] i.e. with an initial hematocrit up to 5% flowing at 1, 2 or 3 mL h^-1^. In addition, we have compared the performance of pristine with surface modified membranes, using commercially available BPS membranes as a benchmark. Finally, non-specific binding of circulating DNA was evaluated for each of the membranes via fluorometric quantification with Qubit™ analysis in order to assess their suitability in use in clinical applications.

## 2 Experimental

### 2.1 Materials

Two different grades of polylactic acid were chosen for this work: Resomer® L 209 S (Poly-Left-Lactic-Acid, PLLA) for TIPS and DIPS membranes and Natureworks® 2002D (PLA) for ES membranes. PMMA sheets at different thickness were produced by Clarex, Nitto Jushi Kogyo Co. Ltd., JP and supplied by Weatherall Ltd., UK. Hexamethylenediamine (HMDA), methanol (MeOH), 1,4 dioxane, chloroforms (TCM), sodium hydroxide (NaOH) and acetone (Ac) were purchased from Sigma Aldrich, Munich, Germany. 37% Hydrochloric Acid (HCl) solution was purchased from Carlo Erba Reagents. Blood samples were collected from The Scottish National Blood Transfusion Service, Jack Copland Centre, Edinburgh, UK. TIPS, DIPS and ES PLA membranes were compared with commercially available membranes Vivid™ GF (Vivid A) and Vivid™ GX (Vivid B) from Pall. All the reactants were ACS grade (purity > 99%).

### 2.2 Thermally Induced Phase Separation membrane preparation

TIPS membranes were prepared by dissolving PLLA (4 wt%) in an 87/13 w/w dioxane-water mixture at 120 °C. The solution, kept above 60 °C to ensure homogeneity, was hot poured in a cylindrical high-density polyethylene (HDPE) mold. The mold filled with the polymer solution was first immersed for 10 minutes in a 0 °C bath and then moved to a -20 °C bath for 20 minutes in order to allow the complete freezing of the as-obtained structure. Successively, the foams were extracted from the mold and subjected to a washing step in distilled water. Finally, the foams were dried under vacuum for 24 h to ensure the complete removal of any solvent trace. The so obtained membranes were then cut to obtain slices 1 mm thick.

### 2.3 Diffusion Induced Phase Separation membrane preparation

DIPS membranes were produced by dissolving PLLA (8 wt%) in neat dioxane at 120°C. A stainless steel slide was immersed into the solution, cooled at 30°C and then pulled out at a constant speed of 3 cm min^-1^ (dip coating) to ensure a proper final average membrane thickness of 120 µm, based on previous work [17]. The coated slide was immerged for 5 minutes into an 87:13 w:w dioxane-water coagulation bath (in order to avoid this so called “skin effect”), and then it was immerged into a pure water bath for other 5 minutes to ensure a complete phase separation. The temperature of both coagulation baths temperature was fixed at 30 °C.

### 2.4 Electrospun membrane preparation

The polymeric solution for electrospinning was obtained by adding PLA (10 wt%) to 20 mL of TCM:Ac (2:1 v:v), and completely dissolved by stirring overnight at room temperature to obtain homogeneous PLA/TCM:Ac solution [26]. A semi-industrial electrospinning equipment (NF-103, MECC CO., LTD., Japan) was used to prepare PLA nanofibers membranes. The polymeric solution was filled in a 10-mL syringe fitted with a 19 gauge stainless steel needle. The electrospinning was then performed using the following constant parameters: flow rate, 1 mL h^-1^; distance between the needle tip and the collector, 13 cm; supplied high voltage, 15 kV; temperature, 25 °C and relative humidity, 40%. The nanofibers obtained were collected on a grounded rotary drum (diameter = 10 cm, speed = 10 rpm) wrapped in an aluminum foil for 180 min in order to obtain membranes approximately 100 μm thick. The collected PLA ES membranes were subsequently dried for at least 2 days under fume hood in order to remove any residual solvents.

### 2.5 Membrane surface modification

All the membranes were surface modified through a solution of methanol:water (3:7 v:v) and HMDA (1% w:v). The solution was directly pipetted onto each side of the PLA membranes. A 12-hour drying period at room temperature in a fume cupboard followed.

#### Morphological analysis

The morphology of the nanofiber mats was evaluated by scanning electron microscopy, (Phenom ProX, Phenom-World, The Netherlands). The samples were attached on an aluminum stub using an adhesive carbon tape.

### 2.6 Fiber diameter and pore size distribution

Fiber diameter distribution of ES membranes and pore size distribution of TIPS and DIPS membranes were determined using ImageJ as an image processing software. In particular, the plugin DiameterJ is able to analyze an image obtained by SEM and to find the diameter of nanofibers at every pixel along a fiber axis. The software produces a histogram of these diameters and summarizes statistics such as mean fiber diameter [26].

### 2.7 pH stability test

The hydrolysis of the ES mats (10 mm × 40 mm × 100 µm) was performed in 10 mL of solutions (pH 1.0, pH 7.0 and pH 13.0) at 25 °C up to 4 hours. At fixed times, i.e. 0.5, 1, 2 and 4 hours, the membranes were washed thoroughly with distilled water at room temperature, followed by drying under chemical hood for 1 day. After drying, the samples were weighed (m_dry_).

### 2.8 FTIR-ATR analysis

The chemical surface properties of the samples were assessed by spectroscopic analysis. FTIR-ATR analysis was carried out by using a PerkinElmer FTIR/NIR Spectrum 400 spectrophotometer; the spectra were recorded in the range 4000–400 cm^-1^.

### 2.9 Water Contact Angle measurements

Static contact angles were measured on all the samples by using distilled water as fluids with an FTA 1000 (First Ten Ångstroms, UK) instrument. More in detail, 4 μL of distilled water were dropped on the membranes. Images of the water on the surface were taken at a time of 10 s. At least 7 spots of each composite nanofiber mat were tested and the average value was taken.

### 2.10 PMMA device preparation and membranes integration

The 4 PMMA layers were 7.5 cm x 3.2 cm with different thickness selected on their function. The upper (L1) and lower (L4) cover were 2 mm while the inlet channel layer (L2) was 1 mm. The layer containing the membrane holder (L3) were 0.5 mm thick with an engraving of 0.1 mm for DIPS membrane and ES 1L membrane. The same layer was 2 mm thick and 1 mm engraved for TIPS membrane and for ES 10L membranes. The entire membrane holder diameter was 7 mm divided as follow: 5.5 mm cut through and 1.5 mm engraved for ensure the correct adhesion of the membrane and avoid flow leaking to the outlet side due to membrane bypass. The inlet and outlet channels in second and fourth layer were engraved while a circle (4.5 mm diameter) at the end of the inlet channel was cut through as input for the membrane. Membranes and PMMA layers with different thickness were cut using a CO_2_ laser cutter (Epilog Mini 18, Epilog, USA) equipped with a 2 in. lens (101.6 μm beam spot). Residual dust was removed with clean-room tissue wetted with ethanol. Layers were aligned in specific custom-made plates endowed with metallic pins; therefore, individual layers were featured with corresponding alignment holes. The membranes were positioned in the proper layer and then ethanol was spread between PMMA layers with a pipette. The bonding was carried out at 70 °C and 100 bar for 5 minutes in a Carver laboratory press [34].

### 2.11 Microfluidic Device testing

A WPI Al300-220 syringe pump loaded with a 10 mL syringe was used to impose a constant flow through the membranes. Plasma-diluted human blood with initial Hct (Hct_in_) equal to 1%, 3% and 5% were used for this test. All kinds of functionalized and unfunctionalized membranes were tested. The blood flow rates were 1, 2 and 3 mL h^-1^ which corresponds to tested output flow rates from existing HBPS system [33]. Measurements were carried out at three volume points of extracted plasma i.e. 0.5 mL, 1 mL and 2 mL. LabSmith Pressure Sensor was used to measure the blood pressure upstream to the membranes.

Hematocrit (Hct) was measured through an hemoanalyzer (ActDiff2, Beckman Coulter).

Separated plasma was collected and analyzed in aliquots at different volume points (*V*_*i*_) i.e. 0.5 mL, 1 mL and 2 mL. The cumulative Hct (%) was calculated according to the following Equation (1):

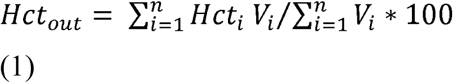

Where *Hct*_*i*_ is the hematrocit evaluated for the i-th sample; *Hct*_*in*_ is the Hct of the feeding blood. The plasma purity [%] was evaluated according to Equation (2):

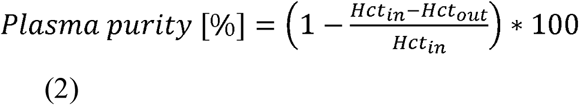

The extent of hemolysis in plasma samples was determined through UV-Vis spectroscopic measurement according to Cripps method. In particular, Cripps number was calculated according to the Equation (3) [35]:

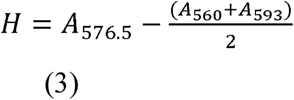

where A_560_, A_576.5_ and A_593_ are the absorbance values at 560 nm, 576.5 nm and 593 nm, respectively. Before UV-Vis spectroscopy, each plasma sample was centrifuged for 10 min at 1,600 g and only the supernatant absorbance was measured in order to ensure to count only free hemoglobin present in the sample. Absorbance was measured using Jenway 7315 Spectrophotometer in 500-630 nm wavelength range at 1 nm interval.

In order to report hemolysis values only due to the cell damage resulting from the microfluidic membranes (H_corrected_), the H value of the centrifuged feed (H_control_) was subtracted to H value of the extracted volume (H_out_) according to Equation (4):

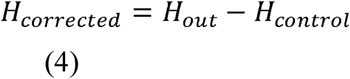

In order to correlate the Cripps number with a concentration of Hgb in plasma, a series of centrifuged plasma containing 0.1 g dL^-1^, to 0.3□g dL^-1^ of Hgb (measured with the hemoanalyzer) were used to obtain a calibration curve, correlating the H value and the hemoglobin concentration. In the concentration range herein investigated, the calibration curve was found to be a line with an angular coefficient of 2.17 up to a H value of 0.33. For higher H value, the calibration curve still remained a line but with a slope of 3.17. In the case of Hgb concentrations higher than 0.3 g dL^-1^, the results from the hemoanalyzer were considered.

Finally, the cumulative Free Hgb (g dL^-1^) concentration was calculated according to the following Equation (5):

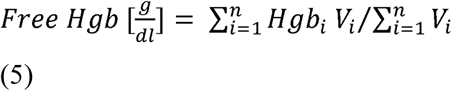

Where *Hgb*_*i*_ is the hemoglobin concentration evaluated for the i-th sample;

The score index calculation was evaluated on the basis of the two following arbitrary parameters: The parameter related with the Hgb concentration (K_Hgb_) was determined according to Equation (6):

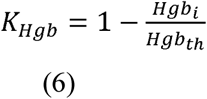

Where, Hgb_th_ is the Hgb threshold value chosen i.e. 0.15 g dL^-1^. Note that if Hgb_i_ > Hgb_th_ then K_Hgb_ < 0 and these values appears in the heat map as a black square.

The parameter related with the plasma purity (K_RBC_) was determined according to Equation (7):

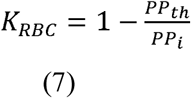

Where, PP_i_ is the plasma purity evaluated for the i-th sample and PP_th_ is the plasma purity threshold value chosen i.e. 85 %. Note that if PP_i_ < PP_th_ then K_RBC_ < 0 and these values appears in the heat map as a black square.

The positive values of K_Hgb_ and K_RBC_ were normalized in the 0-1 range and called nK_Hgb_ and nK_RBC._ The score index calculation (SIC) was then evaluated according to the following Equation (8):

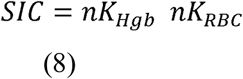

### 2.12 DNA binding

In order to assess the binding capacity of the membranes the mixture of fragmented T47-D cell line DNA (∼160bp), mimicing circulating cell-free DNA fragmentation, was used. DNA solution was prepared containing ∼1 ng µL^-1^ DNA in molecular grade water. To prepare spin columns silica membranes were removed from EconoSpin columns and replaced with ES membranes.[36] The columns were loaded with 50 µL of DNA solution and incubated at room temperature for 30 minutes. After the incubation period, samples were centrifuged at 13,400 rpm for 2 minutes to pull solution through the membranes. DNA concentration within the filtered product was measured using a Qubit dsDNA (double-stranded DNA) High Sensitivity Assay Kit – by Qubit Fluorometric Quantification. A minimum of three membranes treated under the same conditions were measured.

### 2.13 Statistical analysis

Statistical analyses of the data were performed through ANOVA by using the GraphPad Prism version 8 software (GraphPad Software Inc., La Jolla, CA, USA). Significant differences among mean values, where applicable, were determined by the means of ANOVA. In all cases, a value of p<0.05 was considered statistically significant.

## 3 Result and Discussion

In an attempt to achieve rapid and effective blood plasma separation, compatible with Lab-on-Chip or in-line processing, we have developed an hybrid strategy, combining HBPS published previously [33], and a MFU (Figure 1A).

**Figure 1.**
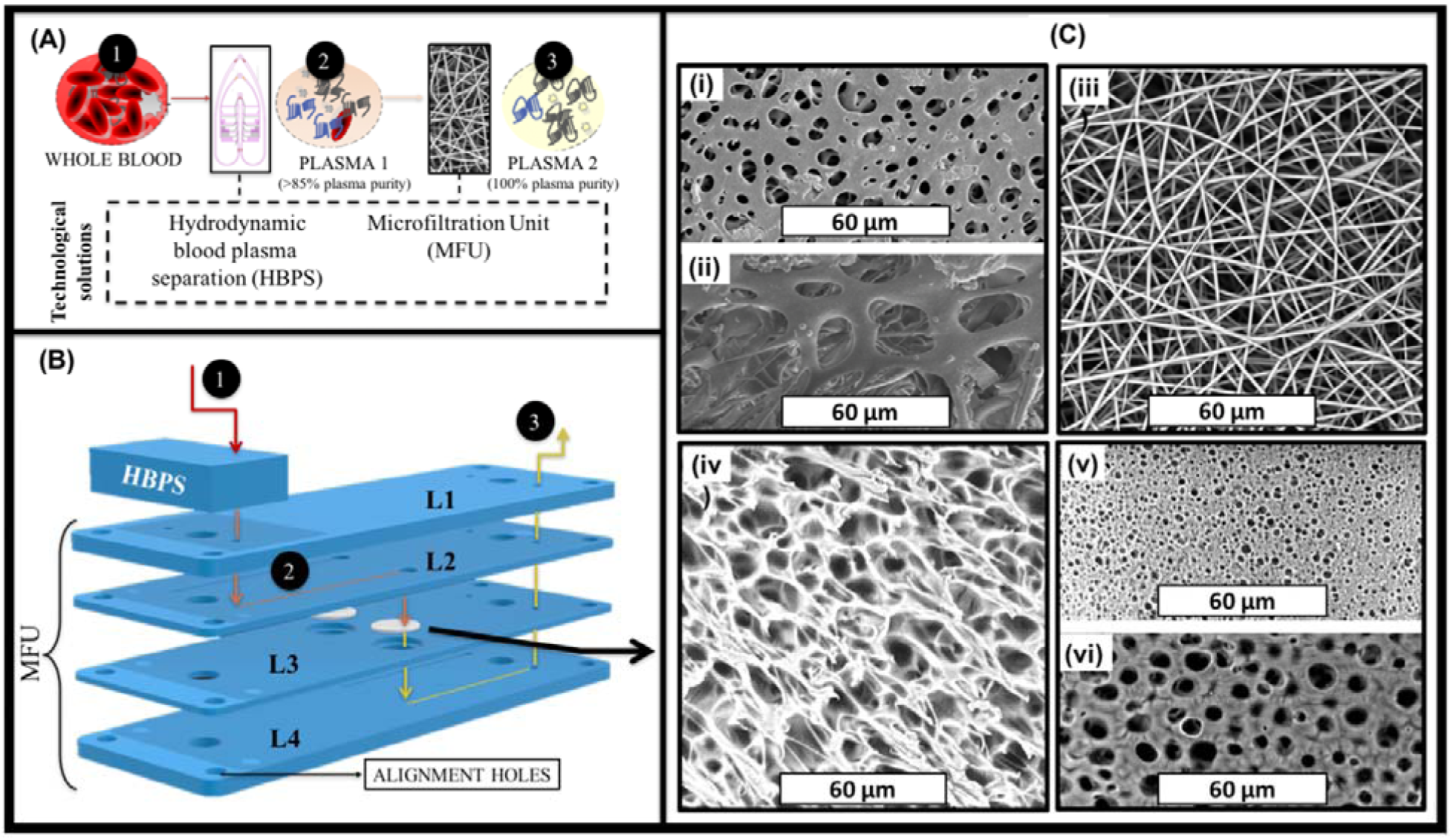
A) Schematic representation of the technical solution for a continuous hybrid blood plasma separation proposed in this work. B) Schematic of the microfluidic device for a continuous RBCs separation proposed in this work. It is composed by an Hydrodynamic blood plasma separation (HBPS) chip and by a Microfiltration unit (MFU). C) SEM images of (i) back side and (ii) front side of Vivid ® membrane; (iii) ES membrane; (iv) TIPS membrane; (v) Back side and (vi) Front side of DIPS membrane.

This two-step strategy should allow the simultaneous achievement of high yield (via the HBPS device) and high purity (via the collection of residual blood cells from the HBPS device on the MFU) in the milliliter range blood plasma separation. In fact, HBPS performance decreases upon increasing plasma yields [9–13]. Combining in the same BPS system hydrodynamic and membrane separation may permit removing the residual RBCs due to an increase of the plasma yields.

Accomplishing high performance separation is not straightforward. Membranes should meet challenging requirements such as low pressure drop and high plasma purity.

In this respect, we have evaluated the use of PLA-based membranes prepared by TIPS, DIPS and ES for the removal of residual red blood cells following HBPS. The performance of these PLA engineered membranes were compared to two commercially available membranes for RBC removal i.e. Vivid™ GF (Vivid A) and Vivid™ GX (Vivid B). As shown in Figure 1B, the membranes were sandwiched in a microfluidic device denoted as MFU and placed in-line after the HBPS device. In this two-step approach, the cells in whole blood (1) are partially separated from the plasma by the HBPS unit (to > 85% plasma purity) (2). They, then, pass through the cover layer (L1) into the L2 inlet channel layer. On the other hand, the partially purified plasma passes through the membrane in L3 into the outlet channel in L4, ready for collection (3).

In the case of both ES and surface-modified ES (m-ES) membranes, 1 layer (ES 1L or m-ES 1L) or 10 layers (ES 10L or m-ES 10L) were placed in the membrane holder, in order to evaluate the effect of different membrane thickness.

For membrane-based cell separation, pore morphology and the size of a membrane are two crucial factors. From the SEM images shown in Figure 1C, it can be seen that a range of different morphologies were obtained depending on which of the three distinct membrane production techniques had been employed i.e. DIPS, TIPS and electrospinning. In addition, Figures 1Ci and ii show the morphologies of the back and front sides of the commercially available Vivid A membrane, (the morphology of Vivid B is very similar to Vivid A and so the images are not included for the sake of brevity). The back side of Vivid membranes (Figure 1Ci) were characterized by pores ranging from 1 up to 9 micrometers in diameter with a mean of 2.84 µm whereas the front side of Vivid membranes (Figure 1Cii) were characterized by pores ranging from 7 up to 29 µm in diameter with a mean of 21.12 µm. From Figure 1Ciii, it was determined that all the fibers in the ES membranes were in the nanoscale range (mean fiber diameter 0.83 µm ± 0.11 µm), randomly oriented and smooth, without any presence of beads. Image processing analysis revealed that the mean pore diameter of this system was 3.53 µm ± 1.67 µm. Figure 1Civ shows the TIPS membrane morphology which was characterized by pores with a mean pore diameter of 3.31 µm ± 0.81 µm and a pore wall of approximately 1 µm. Figures 1Cv and vi show, the morphologies of the back (or bath side) and front sides (slide side) of the DIPS membrane, respectively. DIPS membranes presented considerable differences in pore morphology between the two sides with the mean pore diameter of the front side (in contact with the stainless steel slide) being 1.02 µm ± 0.19 µm while that for the back side (the side immersed in the water/dioxane solution) was 3.05 µm ± 0.36 µm. This asymmetry in pore morphology can be ascribed to the water diffusion kinetic into the polymeric solution. In particular, during desiccation, the back side (directly exposed to air) is characterized by faster water removal compared to the front side (in contact with stainless steel). For this reason, the external membrane surface is more likely to be closed, as faster water removal could re-establish a single-phase condition [17].

### 3.1 Comparative study on PLA membranes degradation at extreme pH values

Chemical based surface modification protocols may be used to provide various functionalities to the membranes. In order to compare the effects of surface modification chemistries, the membranes were subjected to extreme pHs (acidic and basic treatment) for up to 4 hours, as shown in Figure 2.

**Figure 2.**
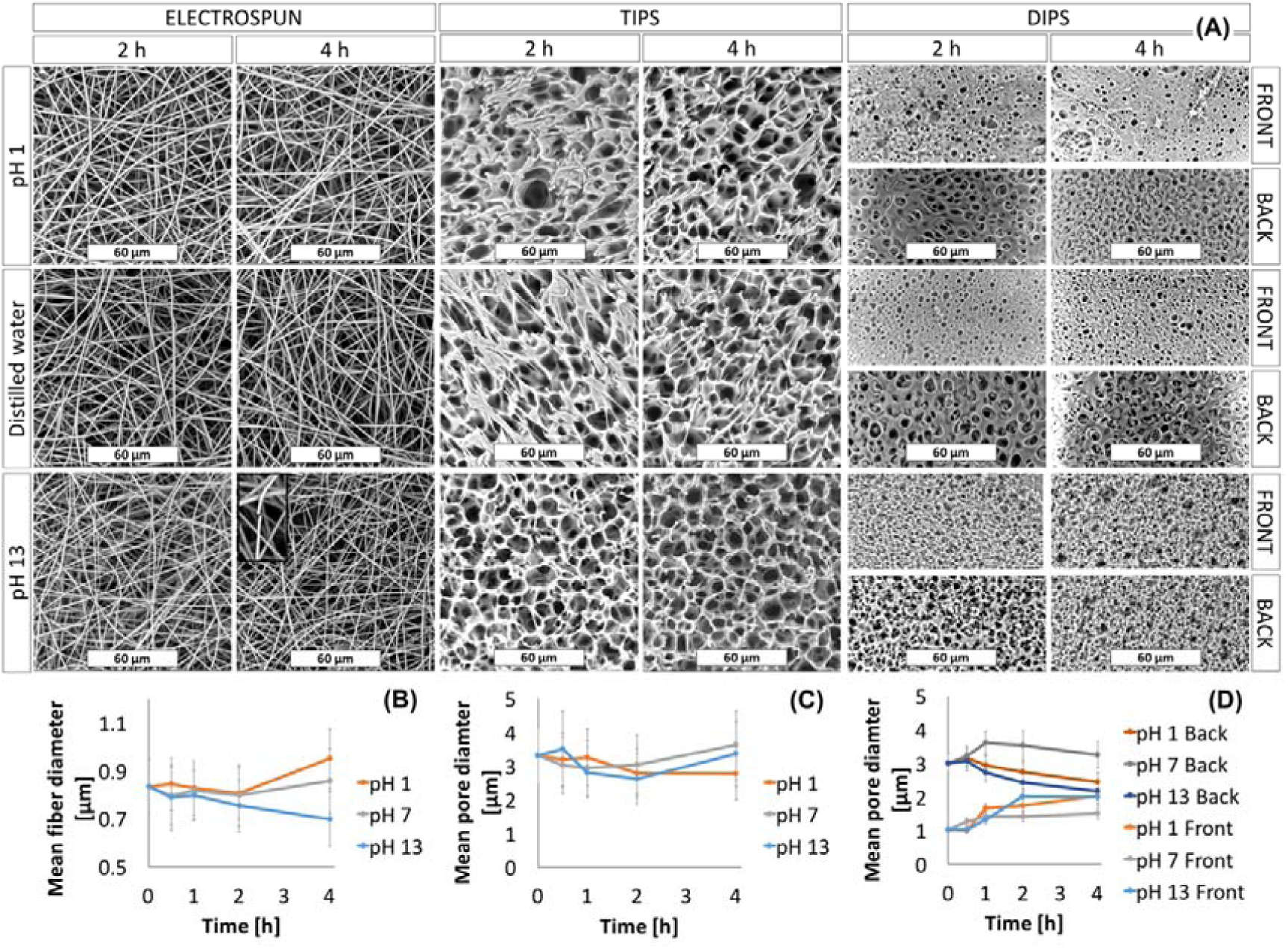
(A) SEM images of membranes immersed for 2 and 4 hours in different medium as a function of pH; (B) Mean fiber diameter of ES membranes as a function of time and pH; Mean pore diameter of (C) TIPS and (D) DIPS membranes as a function of time and pH.

In particular, the hydrolytic degradation behavior of the membranes was assessed in three different pH, pH 1, distilled water and at pH 13.The morphology of the membranes at 25 °C for two time points (2 h and 4 h), are shown in Figure 2A while in Figure 2B, 2C and 2D the mean fiber diameter of the ES mats and the mean pore diameter of the ES, TIPS and DIPS membranes, respectively, are reported.

ES membranes immersed in acidic medium and distilled water for up to 4 hours did not show any significant change in their morphology. In fact, only a slight increase in the fiber diameter (Figure 2B) in acidic medium after 4 hours was observed. On the other hand, ES membranes immersed in the medium at pH 13 for 4 hours exhibited a considerable number of broken fibers (see the inset) and a slight reduction in the mean fiber diameter that did not affect the average pore size.

Similarly, TIPS and DIPS membranes showed a more pronounced morphology change in the basic medium. Although the mean pore diameter of the TIPS remained almost constant (Figure 2C), it can be clearly seen that, at pH 13, the edge of the pores had become partially eroded, particularly after 4 hours.

DIPS membranes immersed in all kinds of medium show a more significant change in mean pore diameter as a function of time. In Figure 2D it is possible to observe that the pore size of the front side gradually increased while that of the back side gradually decreased, in particular for membranes immersed in pH 1 and pH 13 solutions. As can be seen in Figure 2A, when the polymer matrix was immersed in a medium at pH 1, the back side seemed to swell, thus inducing a reduction in the pore size, while the edges of the pores on the front side partially eroded with time, thereby slightly increasing the pore diameter. A similar phenomenon was observed for the membranes immersed in distilled water, although less pronounced. In contrast, DIPS membranes immersed in a medium at pH 13 seem to be affected by deeper erosion. In fact, after 4 hours both side of the membranes show a similar morphology likely due to the degradation of the membrane external layers (Figure 2A).

The above observations can be explained by considering the mechanism of hydrolysis of PLA, which is primarily determined by the amount, presence and location of water molecules. Recent studies correlated the differences in the degradation behavior of PLA to the hydrophilicity of this polymer matrix exposed in media at different pH [37]. PLA exposed to basic media (pH 13) maintains its non-polar (hydrophobic) character, thus reducing its absorption capacity of water. Therefore, water cannot permeate the polymer matrix thus causing superficial degradation. Differently, in acid medium (pH 1) PLA molecular chains change their character from hydrophobic to hydrophilic thus allowing PLA swelling and the bulk degradation of the samples [19]. In both cases, during PLA degradation, lactic acid oligomers, lactide and lactic acid are formed [38]. In conclusion, the PLA membrane durability in the acid medium is very similar to that in the neutral medium and higher than that in the alkaline environment. Probably the PLA bulk degradation due to acidic medium is slower than surface degradation due to basic environment thus affecting the membranes morphology less than the basic environment during the 4 hours of the test time.

### 3.2 Wettability and surface functionalization of PLA-based porous or fibrous membranes

In order to accomplish high performance separation, membranes should meet challenging requirements such as low pressure drop that can be reduced by modifying their wettability.

Several approaches have been investigated in order to enhance the wettability of PLA-based porous or fibrous membranes in both dry and wet chemistry [29–31]. In this work, an aminolysis surface treatment via hexamethylenediamine (HDMA) was carried out. SEM images of PLA-based membranes after aminolysis are shown in Figure 3.

**Figure 3.**
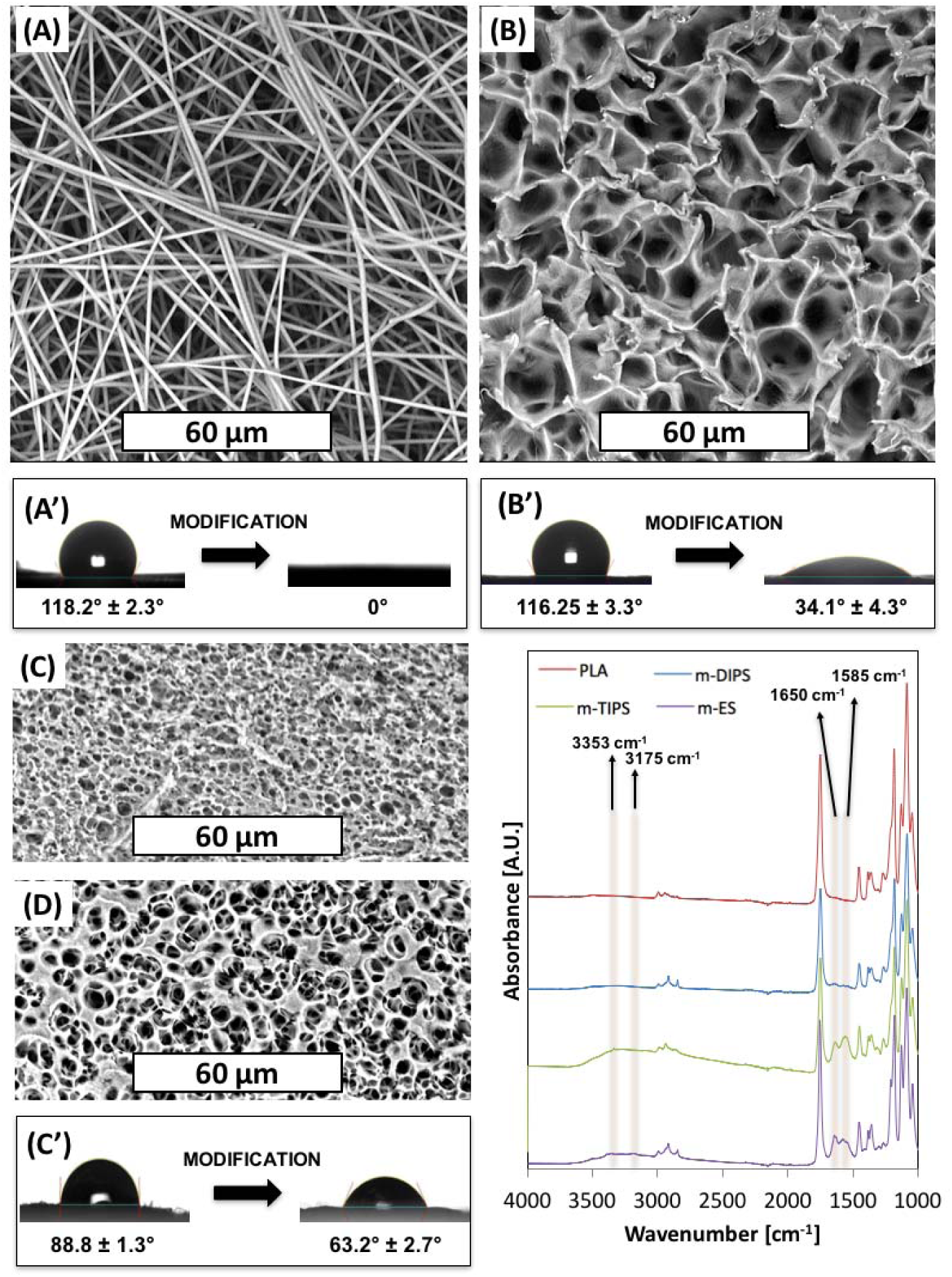
SEM images of (A) ES membrane; (B) TIPS membrane; (C) Front side and (D) back side of DIPS membrane after aminolysis. Effect of surface modification on WCA of (A’) ES membrane; (B’) TIPS membrane; (C’) DIPS membranes; (E) FTIR-ATR measurements of PLA and membranes after aminolysis.

Morphological inspection clearly reveals that the surface treatment eroded the porous structure of all membranes. In particular, from Figure 3A, for the m-ES, it is possible to observe several fractured fibers, as already reported by others [39]. Furthermore, it can be seen that some fibers are attached to each other in bundles of 2 to 6. The reason for this phenomenon may be due to the increased interaction now possible following surface modification. Similar to the treatment in basic medium, the TIPS membrane (Figure 3B) showed erosion of the pore edges while the DIPS membrane showed partial erosion of the superficial layer of the membrane.

In Figure 3A’C’ the effect of surface modification on the water contact angles are reported. Unmodified ES, TIPS and DIPS membranes showed WCA equal to 118°, 116° and 89°, respectively. It is well known that the water contact angle is dependent on surface chemical functional groups, porosity and surface roughness. Since the polymer matrix is the same in all cases, this result can be reasonably ascribed to the higher porosity of both the TIPS and ES membranes with respect to the DIPS membranes. In fact, porosity allows the material to entrap air when the surface is immersed in water, thus enhancing the hydrophobicity of the PLA membranes [19]. At the same time, m-ES and m-TIPS membranes showed the highest WCA reduction, i.e. 100% and 71% respectively, while WCA decrease of m-DIPS membranes was only 29%. This result can be likely ascribed to the higher specific surface of ES and TIPS membranes that permits exposing more amine moieties to water drops than DIPS membranes.

Figure 3E shows the FTIR-ATR spectra of neat PLA and amino-modified ES, TIPS and DIPS membranes. The absorption spectra of PLA showed typical peaks such as the C=O stretch at 1747□cm^-1^, the C-O stretch at 1180, 1129 and 1083□cm^-1^ and the OH bend at 1044□cm^-1^[26]. The presence of amine functional groups on PLA can be observed by the vibrations of N–H (asymmetric stretching), N–H (symmetric stretching), the overtone of N–H deformation and N–H primary amine deformation, which give rise to peaks at 3353, 3262, 3175 and 1585 cm^-1^, respectively [40]. Although absorption in the range 3000 - 3500 cm^-1^ in amino-modified membranes is barely visible due to the vibration of amide groups that lie in the same regions and to the presumable presence of water due to the enhanced wettability, the peak at 1585 cm^-1^ is clearly visible in particular for m-TIPS and m-ES membranes. Moreover, after aminolysis of ES, TIPS and, to a lesser extent, DIPS membranes showed another new peak at 1650 cm^−1^ which corresponded to amide I stretch [41]. The intensities of the amine-related peaks, including the peak related to the primary amide, decrease progressively for ES, TIPS and DIPS membranes and this is consistent with the WCA results of amino-modified membranes showing that the reactive amine density is highest at the highest specific surface exposed by the membrane [40].

### 3.3 Membrane performance: Plasma purity

In order to evaluate the membranes performance, they were included in a microfluidic device denominated MFU. The MFU design is shown in Figure 1B while top and cross section photographs of the device are available as Supporting Information (SI 1).

The membrane performance was first evaluated through an investigation of the plasma purity. The purity of the plasma output from the devices utilizing either the ES membrane (amino-modified or not) or one of the two commercially available membranes for RBCs separation (Vivid A and B) was determined by comparing the inlet Hct (Hct_in_) of the blood sample with the outlet Hct (Hct_out_) according to Equation 2 as a function of different flow rates and extract plasma volume.

Similar experiments performed using both the surface modified and unmodified TIPS and DIPS membranes resulted in clogging (reaching a pressure higher than 300 kPa) before 0.5 mL of plasma had been extracted. The results have been compiled into a three-color heat map and shown in Figure 4A.

**Figure 4.**
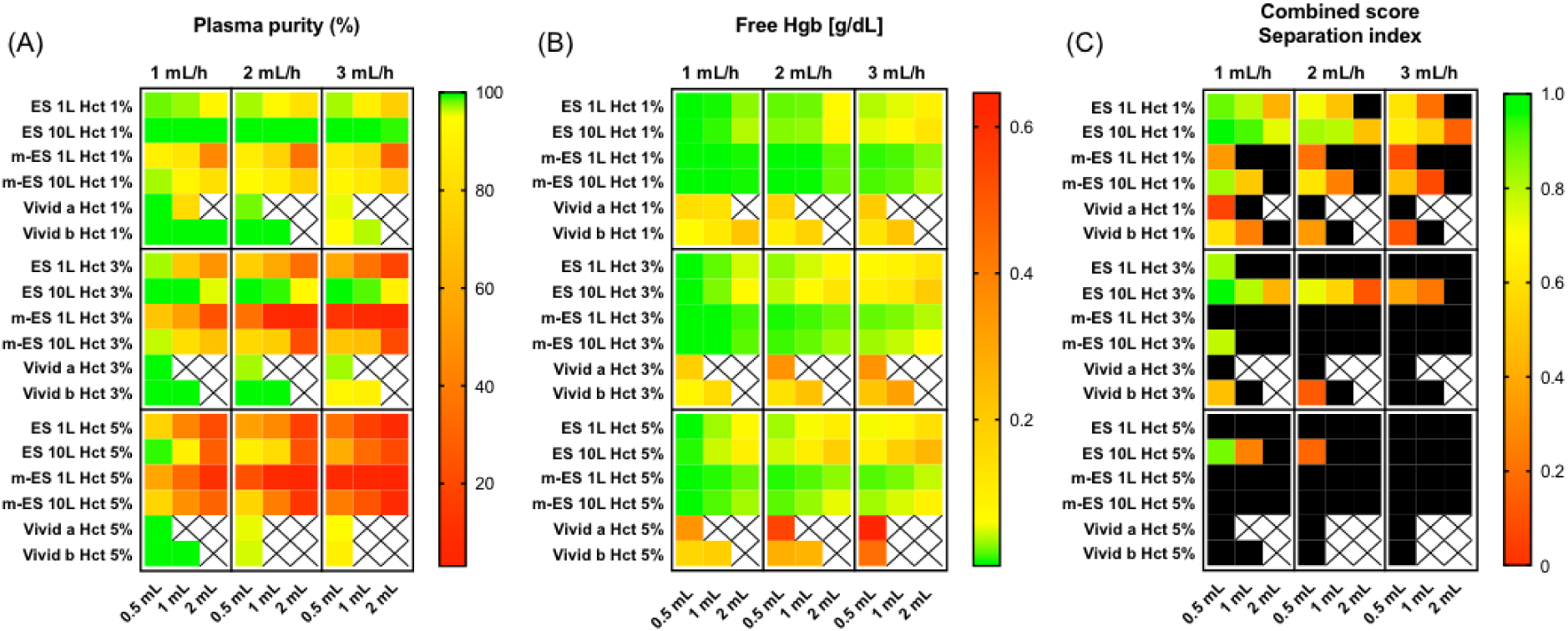
Electrospun Membrane Low Hematocrit Separation Performance. A) Plasma purity is represented by a triple color gradient from green (100%, corresponding to no cell detected) to red (0%) as a function of Hctin for three different treated blood volume (0.5, 1 and 2 mL) and blood flow rate (1, 2 and 3 mL h-1), organized in quadrants for clarity. The baseline value is set at 95% and represented in yellow, to enable rapid reading and selection (green corresponding to plasma purity > 95%). Crosses indicate sets of parameters were data was not acquired, because the devices could not handle high pressures. B) Free hemoglobin concentration (g dL-1) as a function of Hctin for three different treated blood volume (0.5, 1 and 2 mL) and blood flow rate (1, 2 and 3 mL h-1) and represented as a triple gradient from green, (0 g dL-1) to red (6 g dL-1). The baseline value is set as 0.06 g dL-1 (yellow). C) The combined score or separation index is calculated from the plasma purity (%) and free Hgb values. A score of 1 indicates the maximum score for 100% plasma purity and 0 g dL-1 free Hgb. The score index calculation was evaluated according to Equation 8. Here, the double threshold is set at 0.15 g dL-1 (Maximum value) and 85% plasma purity (Minimum value) at which point, the score appears in the heat map as a black square. The baseline value is set at 0.5 arbitrary units. The threshold can be changed to enable the user to exclude underperforming membranes from the selection (see SI 2 for other threshold values).

The general trend observed is that there is a decrease in plasma purity with increasing flow rate, residual hematocrit and extracted volume. The highest plasma purity was achieved using ES 10L. Employing this device, it was possible to extract up to 2 mL of pure plasma (100% of plasma purity) from plasma with 1% residual Hct_in_ with all the flow rates investigated. For plasma samples containing 3% residual Hct_in_, the plasma purity decreased to 96%, 93%, and 90% using flow rates of 1 mL h^-1^, 2 mL h^-1^ and 3 mL h^-1^, respectively. It is worth noting that 90% of plasma purity due to the MFU fed with 3% of Hct_in_ represents a global plasma purity (i.e. due to both HBPS chip and MFU) higher than 99%. In fact, assuming a whole blood hematocrit of 40%, 3% Hct_in_ can be achieved by an HBPS chip plasma purification of around 92.5%. After a further microfiltration step able to achieve 90% plasma purity, the global plasma purity of the hybrid system would be 99.25%. The worst plasma purity was observed in devices containing m-ES 1L membranes. The commercial membranes compared relatively well at the lowest flow rate, and the lowest residual Hct, but rapidly clogged, and did not yield results beyond this (no data collection represented by squares on heat map), showing limited use at higher volumes.

### 3.4 Membrane performance: Hemolysis

Membrane performance was also evaluated by measuring the free hemoglobin concentration in the extracted plasma due to the RBCs rupture during their passage through the membrane.

The two performance indicators, plasma purity and hemolysis, are needed to reflect on the true performance of the device as a high RBCs removal on its own would not necessarily reflect an overall good performance if the filtrates have high levels of hemoglobin.

In order to assess the amount of hemolysis due to membrane separation, the free hemoglobin concentration was determined according to Equation 5, as described in the ‘materials and methods’ section. The results have been reported in a heat map format (Figure 4B). As expected, the general trend observed is that there is an increase in the free Hgb concentration upon increasing the flow rate, Hct_in_ and the extracted volume. In this case, the best performances were shown by the surface modified (aminolyzed) membranes. Specifically, if 1% Hct_in_ is used as input for the devices incorporating either m-ES 1L and m-ES 10L membranes, the concentration of free hemoglobin in the output samples was lower than 0.04 g dL^-1^. On the contrary, RBCs lysis was found to be particularly pronounced for devices including the commercial porous membranes, Vivid A and Vivid B, at all the flow rates here investigated. For instance, Vivid B membranes showed a concentration of free hemoglobin equal to 0.21 g dL^-1^ after 1 mL of plasma extracted, with a flow rate equal to 3 mL h^-1^.

Upon increasing the Hct_in_ up to 3% and 5% the free Hgb concentration increased for all the system investigated but the trend remained similar (Figure 4B).

### 3.5 Membrane performance: Combined score

It is worth noting that high separation efficiencies can result in high hemolysis value. Both performance indicators need to be balanced, to achieve and overall good separation with maximum plasma purity and minimal hemolysis. For this reason, in Figure 4C we report a combined score calculated using Equation 8 which takes both plasma purity and hemolysis into account. The higher the score, the higher the performance of the membrane. A double threshold can be set, so that systems displaying one or both indicators beyond certain thresholds can be excluded from the selection matrix. In our example in Figure 4C, we have arbitrarily set a threshold at 85% plasma purity and 0.15 g dL^-1^ hemoglobin level. Membranes displaying indicators respectively below or above these levels are represented in black squares. Threshold can be set up to facilitate selection when testing a large number of membranes. Here, the 0.15 g dL^-1^ hemoglobin level corresponds to a threshold for moderate hemolysis according to Snyder et al. [42]. The heat map clearly reveals that the best combined performance was achieved by the ES 10L membranes since they offered the best compromise between the plasma purity and a relatively low hemolysis degree. Combined score separation index heat maps obtained with other thresholds are available in Supporting Information (SI 2).

### 3.6 Membrane pressure drop

It is worth noting that hemolysis is strongly affected by the pressure drop across the membranes because it can stress the RBC membrane. In Figure 5 it is reported the pressure drop across all ES (amino-modified or not) and commercial Vivid membranes as a function of the hemolysis (free hemoglobin concentration). As expected, Figure 5 shows that free Hgb concentration increases upon increasing the maximum pressure drop across the membranes. The Vivid membranes (red dots) always showed higher hemolysis for the same pressure drop for the corresponding ES membranes (black dots).

**Figure 5.**
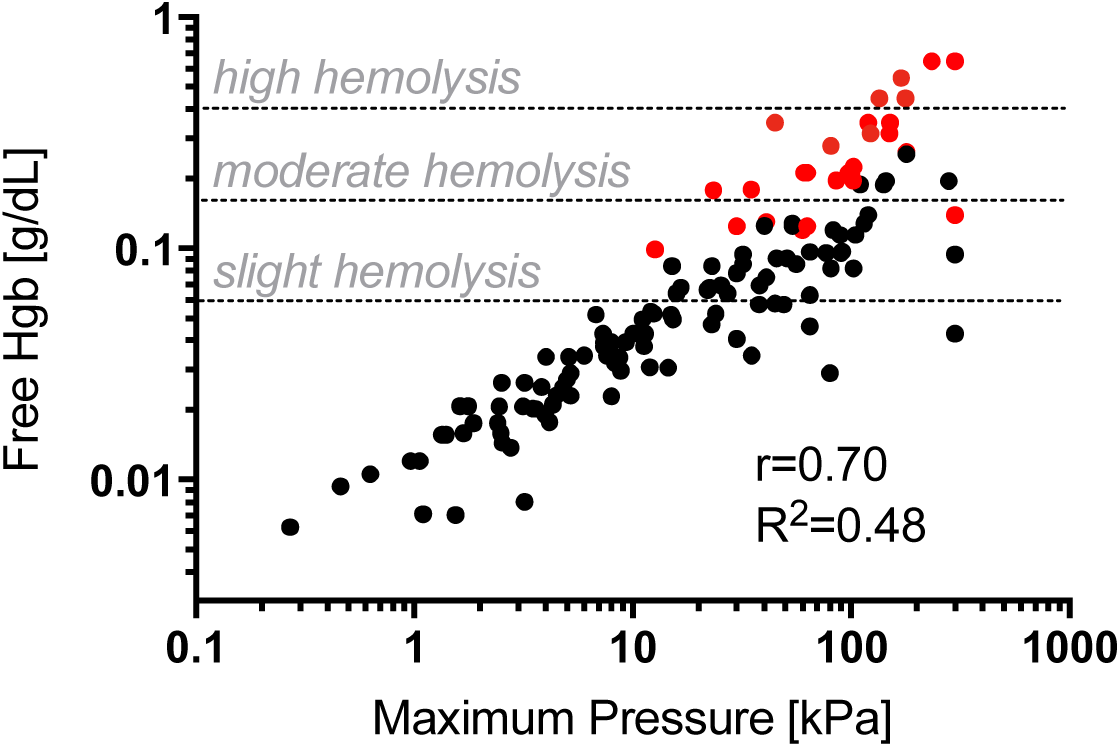
Free hemoglobin represented as a function of maximum pressure recorded. Each point is obtained through the average of at least three independent experiments (n=3). For clarity we have only highlighted one indicator. The commercial membranes plotted in red, clearly show higher pressure drops and resulting hemolysis. We used Snyder scale of Hgb concentrations of 0.060, 0.150, and 0.300 g dL-1 corresponding to thresholds for slight, moderate, and high hemolysis [42]. The slight, moderate and high levels are showed with three horizontal dash lines.

Hemolysis as a result of membrane separation can be attributed to several factors including the shear stress on the RBCs, membrane morphology and the pressure sustained by the cells while in the device. The shear stress on RBCs increases upon increasing the flow rate and this can also affect the pressure on the membrane. The porous structure of membranes can affect both the pressure drop and the damage induced through contact between the cell and the pores. In the Supporting Information (SI 3), the pressure drops values in MFU are reported as a function of the diluted-blood flow rates and the extract plasma volume. As expected, the general trend is that there is an increase in the pressure drop upon increasing the flow rate, Hct_in_ and the extracted volume.

Both commercial membranes, Vivid A and Vivid B, showed the highest pressure drops, although it should be noted that these membranes have been developed for low volume separation from whole blood. On the other hand, in this work, Vivid membranes were used for high blood volume separation but only to purify plasma with minimal residual hematocrit, not whole blood. As expected, ES 10L showed higher pressure drops when compared to ES 1L because of its higher thickness. Regardless of flow rates, the ES membrane pressure drop was 50 - 60% lower than the Vivid membranes at 1% Hct_in_. The pressure drop differences between ES and Vivid membranes can be likely ascribed to the nanofibrous structure of the mats which offers large interconnection degree and, as a consequence, low pressure drop. Moreover, despite their poor separation performance, the aminolyzed membranes exhibited even lower pressure drops due to their enhanced wettability as a result of the chemical functionalization.

The pressure drop differences can partially explain the differences in hemolysis levels between Vivid and ES membranes. It is envisaged that the fibrous morphology of the latter’s PLA-based mats and their superior interconnection degree reduced RBCs lysis during plasma separation. In fact, during contact with the red blood cell, the ES fibers may bend thus reducing the stress on the RBC extracellular membrane and, as a consequence, their lysis.

### 3.7 Non-fouling performance

The cell separation membranes have been designed to be integrated into a complete workflow for circulating DNA extraction. Avoiding DNA adsorption onto the surface of the membrane is therefore crucial for the functioning of the membrane. To investigate the non-fouling properties of the membrane, DNA extraction after permeation of a plasma sample through the membrane placed in a column has been performed.

In Figure 6 the DNA binding test procedure draws and results are reported. In brief, in order to assess this test, silica membranes were removed from commercial spin columns and replaced with the ES membranes (Figure 7A) [36]. The columns were loaded with the DNA solution (1 ng µL^-1^), incubated, and then centrifuged to pull the solution through the membranes (Figure 6B, Materials and Methods).

**Figure 6.**
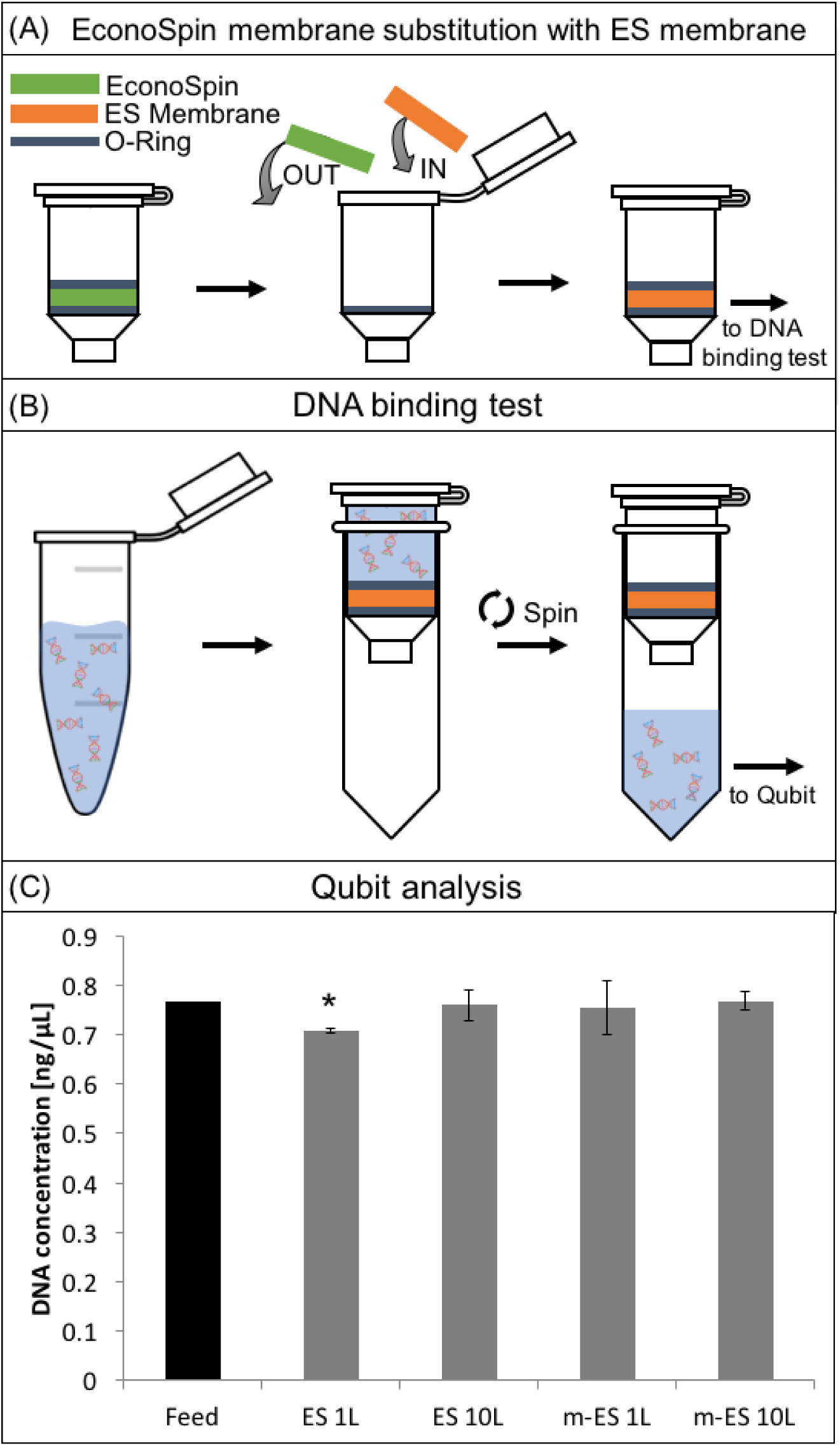
A) Schematic describing the substitution of the membrane in a column device with the engineered electrospun membranes; B) schematic describing the DNA binding test procedure; C) DNA concentration in water before (control) and after permeation trough modified and unmodified electrospun PLA membranes. * Indicates statistically significant differences (p<0.01) when analyzed by ANOVA.

The histogram in Figure 6C represents the concentration of DNA before (positive control) and after the permeation trough ES layers. Results showed that ES 10L and m-ES 1L and m-ES 10L did not significantly (p < 0.01) affect the DNA concentration. Only ES 1L lead to a slight decrease of DNA concentration, within the instrument deviation range. These results highlight that both the unmodified and modified ES PLA-based membranes are low binding and suitable for use in assays where maximum DNA recovery from raw samples is of importance.

## 4 Conclusion

PLA-based membranes prepared by DIPS, TIPS and electrospinning were modified trough aminolysis and characterized. The membranes resistance was tested at extreme pH values for up to 4 hours. Basic media was the most aggressive environment affecting pore morphology. The successful amine functionalization was demonstrated by FTIR-ATR while the subsequent change in wettability of the samples was assessed though WCA analysis. Both modified and unmodified TIPS and DIPS membranes clogged before extracting 0.5 mL of plasma and were excluded from further examination. ES membranes were selected for blood plasma separation tests at low hematocrit levels (1, 3 and 5%), corresponding to output-plasma tested from already existing HBPS systems. Single layers (ES 1L) or stacks of 10 ES membranes were tested in a microfluidic configuration devise. The stack of 10 layers (ES 10L) allowed the collection of up to 2 mL of RBCs-free and low-hemolyzed (free Hgb = 0.078 g dL^-1^) plasma at 1% Hct_in_ and flow rate up to 2 mL h^-1^. With the same membrane and flow rate, upon increasing the Hct_in_ the plasma purity decreased but it was also possible to collect 1 mL of RBCs-free and low-hemolyzed (free Hgb = 0.062 g dL^-1^) with 3% Hct_in_. There is evidence to show that ES membranes decreased the degree of hemolysis when compared to the commercially available Vivid ® membranes. The ES membranes also exhibited lower pressure drops while maintaining comparable plasma purity, thereby showing superior separation performance at low hematocrit levels. We have shown that the PLA-based ES membranes may be suitable for liquid biopsy workflows, where maximum recovery of circulating DNA is crucial. PLA-based ES membranes can be readily integrated into microfluidic workflows and show enhanced separation performance compared to commercial membranes, for the recovery of high purity plasma in the hundreds of microliters to milliliter range.

## Acknowledgements

A.O is funded by a James Watt scholarship.

M.K.K. acknowledges funding from the Engineering and Physical Sciences Research Council, EP/R00398X/1.

F.L. is funded by the European Social Fund (ESF) - A.I.M: Attraction and International Mobility_ AIM1845825 – 1 CUP: B74I18000260001

