## Supporting Info 1, 2, 3. for "Engineered membranes for residual cell trapping on microfluidic blood plasma separation systems. A comparison between porous and nanofibrous membranes"

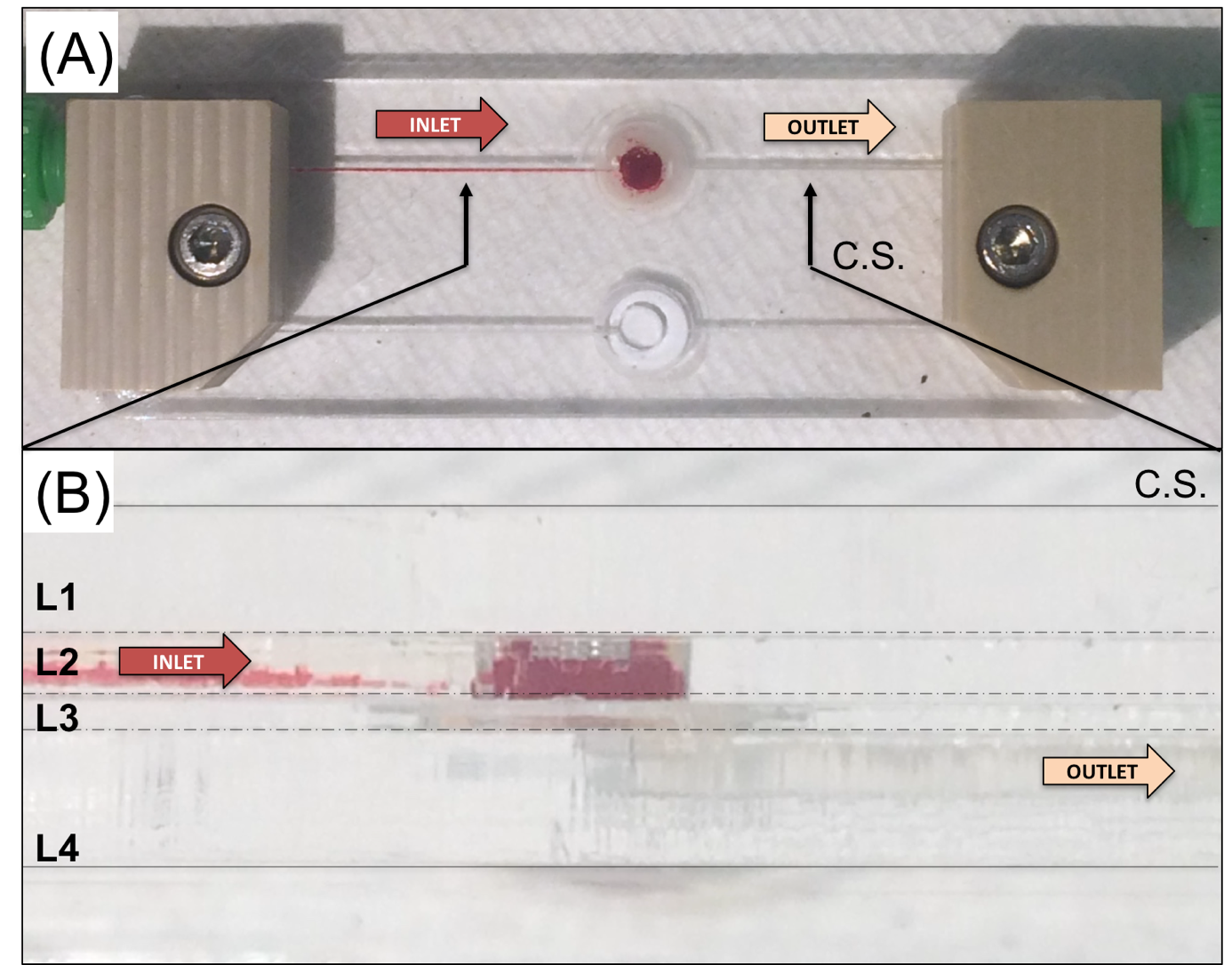

*Supporting Information 1 -Pictures of A) Top view and B) cross section of the assembled microfiltration unit.*

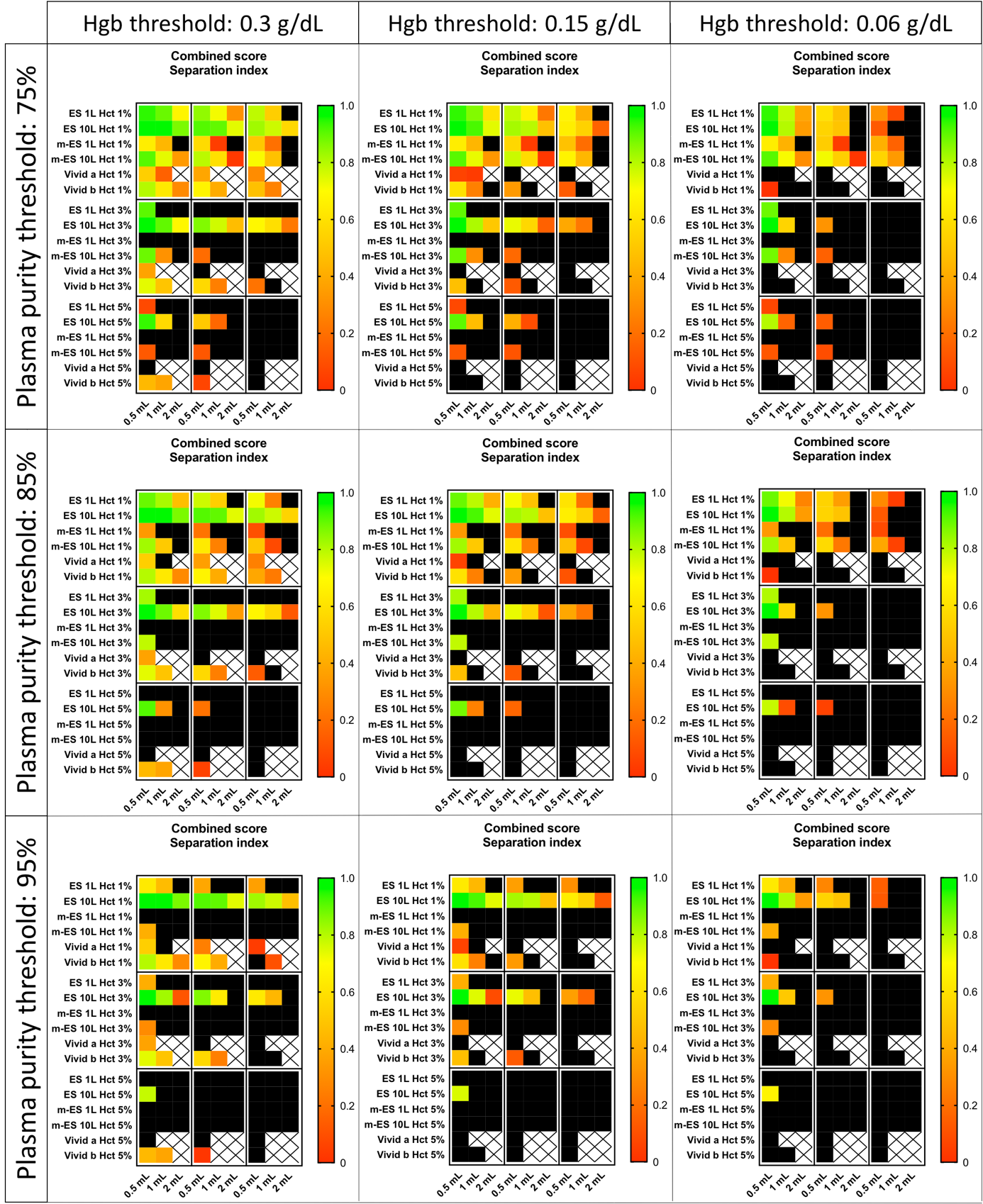

*Supporting Information 2 - Combined score separation index heat maps evaluated according to Equation 8 as a function of different plasma purity and free Hgb threshold values.*

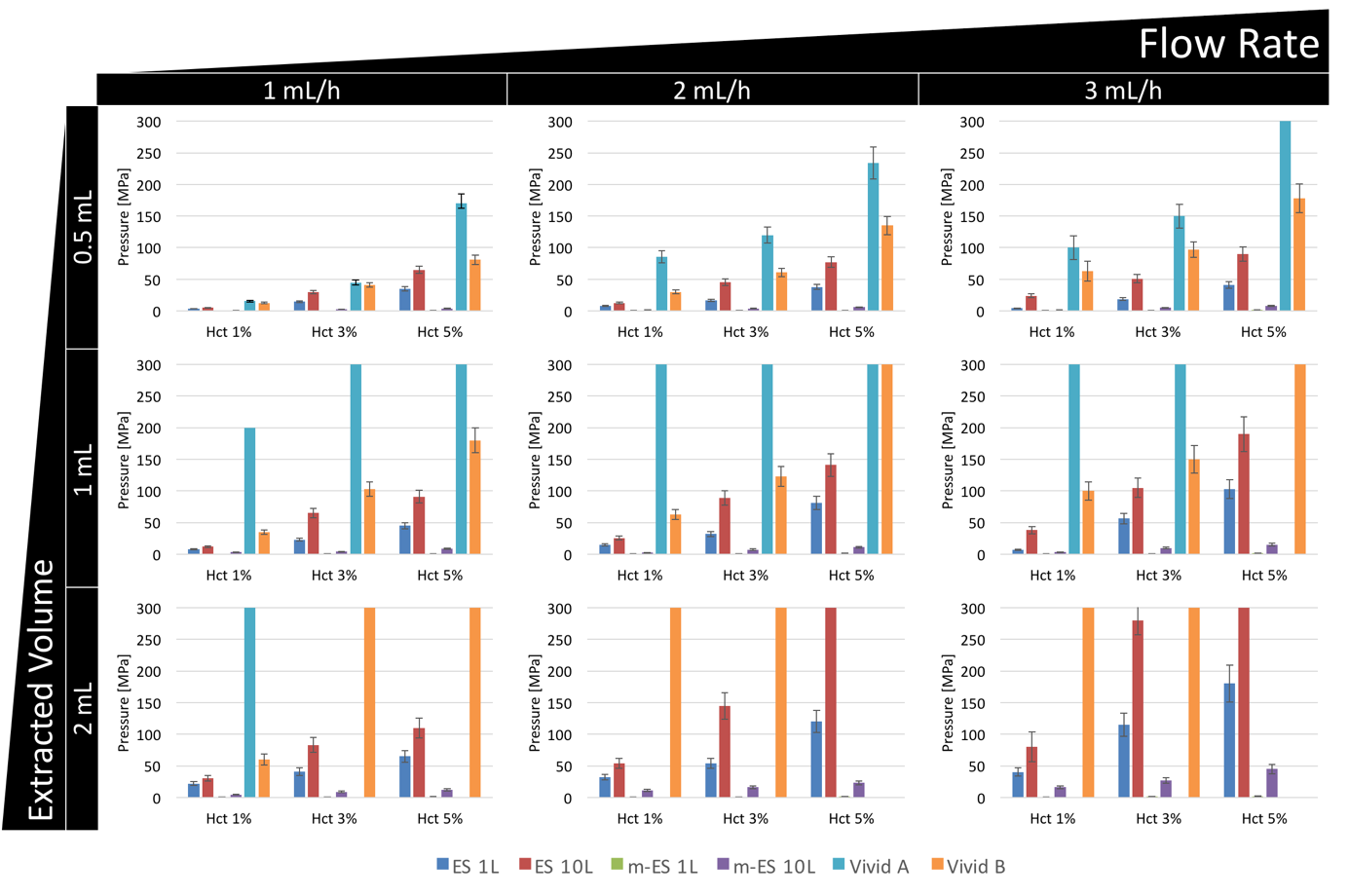

*Supporting Information 3 – Maximum pressure measured in the MFU as a function of the extracted plasma volume, inlet blood flow rate, Hct_in_ and the kind of membrane.*
